# Coinfection of *Streptococcus pneumoniae* reduces airborne transmission of influenza virus

**DOI:** 10.1101/2020.11.10.376442

**Authors:** Karina Mueller Brown, Valerie Le Sage, Andrea J. French, Jennifer E. Jones, Gabriella H. Padovani, Annika J. Avery, Michael M. Myerburg, Stacey Schultz-Cherry, Jason W. Rosch, N. Luisa Hiller, Seema S. Lakdawala

## Abstract

Secondary bacterial infection, especially with *Streptococcus pneumoniae* (Spn), is a common complication in fatal and ICU cases of influenza virus infection. During the H1N1 pandemic of 2009 (H1N1pdm09), there was higher mortality in healthy young adults due to secondary bacterial pneumonia, with Spn being the most frequent bacterial species. Previous studies in mice and ferrets have suggested a synergistic relationship between Spn and influenza viruses. In this study, we used the ferret model to study whether airborne transmission of H1N1pdm09 was influenced by coinfection with two Spn serotypes: type 2 (D39) and type 19F (BHN97). We found that coinfected animals experienced more severe clinical symptoms as well as increased bacterial colonization of the upper respiratory tract. In contrast, we observed that coinfection resulted in reduced airborne transmission of influenza virus. Only 1/3 animals coinfected with D39 transmitted H1N1pdm09 virus to a naïve recipient compared to 3/3 transmission efficiency in animals infected with influenza virus alone. A similar trend was seen in coinfection with BHN97, suggesting that coinfection with Spn reduces influenza virus airborne transmission. The decrease in transmission does not appear to be caused by decreased stability of H1N1pdm09 in expelled droplets in the presence of Spn. Rather, coinfection resulted in decreased viral shedding in the ferret upper respiratory tract. Thus, we conclude that coinfection enhances colonization and airborne transmission of Spn but decreases replication and transmission of H1N1pdm09. Our data points to an asymmetrical relationship between these two pathogens rather than a synergistic one.

**Significance:** Airborne transmission of respiratory viruses is influenced by many host and environmental parameters. The complex interplay between bacterial and viral coinfections on transmission of respiratory viruses has been understudied. We demonstrate that coinfection with *Streptococcus pneumoniae* reduces airborne transmission of influenza A viruses by decreasing viral titers in the upper respiratory tract. Instead of implicating a synergistic relationship between bacteria and virus, our work demonstrates an asymmetric relationship where bacteria benefit from the virus but where the fitness of influenza A viruses is negatively impacted by coinfection. The implications of exploring how microbial communities can influence the fitness of pathogenic organisms is a novel avenue for transmission control of pandemic respiratory viruses.

## Introduction

Acute pneumonia occurs in 30-40% of influenza A virus (IAV) hospitalizations and can be caused by the virus alone or in conjunction with a secondary bacterial infection, most commonly *Staphylococcus aureus* and *Streptococcus pneumoniae* (Spn) (Kalil & Thomas, 2019; Morens, Taubenberger, & Fauci, 2008). During the 2009 H1N1 pandemic (H1N1pdm09) and the 1918 H1N1 pandemic, secondary infection with Spn contributed to severity of disease, especially in young adults (Khardori, 2010; Klein et al., 2016; Louie et al., 2009; Morens et al., 2008). Animal models to study the relationship between IAV and Spn have largely focused on immune responses, disease outcome, and the influence of IAV on Spn colonization (Diavatopoulos et al., 2010; Duvigneau et al., 2016; Metzger & Sun, 2013; Mina, Klugman, Rosch, & Mccullers, 2015; Mina, McCullers, & Klugman, 2014; Nakamura, Davis, & Weiser, 2011). Given the epidemiological importance of viral-bacterial coinfections, there is a critical need for studies that examine how bacterial pathogens influence viral fitness within the host and its transmission to new hosts.

The prevailing model describes a synergism between IAV and Spn with regards to morbidity and mortality. Exacerbated disease in coinfections when compared to single IAV or single Spn infections is supported by epidemiological studies as well as studies in murine and ferret models of respiratory infections. Phenotypic studies in the murine model suggest that heightened disease results from a combination of dysregulation in immune responses, extensive epithelial damage, release of planktonic bacteria from biofilms, as well as increased bacterial colonization and dissemination (Mina & Klugman, 2014; Morens et al., 2008; Rudd, Ashar, Chow, & Teluguakula, 2016). Mechanistic studies indicate that increased bacterial colonization is a consequence of upregulation of bacterial receptors and IAV-induced increases in sialic acid, which serves as a nutrient source for Spn (McCullers & Rehg, 2002; Mina & Klugman, 2014; Siegel, Roche, & Weiser, 2014). Further, enhanced bacterial systemic dissemination is facilitated by virus induced cytokine release, oxidative stress and pneumolysin release (Gonzalez-Juarbe et al., 2020; Mina & Klugman, 2014; A. L. Richard, Siegel, Erikson, & Weiser, 2014). Many of these published studies focused primarily on Spn specific colonization, transmission, and pathogenesis, and do not explore the impact of Spn on IAV binding, replication, or dissemination.

In addition to colonization and disease, a few studies have also investigated the influence of coinfection on contact-dependent or airborne transmission. Contact-dependent transmission studies performed in infant mice clearly show that coinfection with IAV dramatically enhances Spn transmission with the efficiency of IAV being highly variable (Diavatopoulos et al., 2010; Short, Reading, Wang, Diavatopoulos, & Wijburg, 2012). Further, recent studies in the neonatal murine model have demonstrated that Spn infection preceding IAV inoculation can decrease IAV transmission (Ortigoza, Blaser, Zafar, Hammond, & Weiser, 2018). However, adult mice are a suboptimal model for IAV transmission since they are not susceptible to human seasonal influenza virus strains, and transmission of mouse adapted IAV is inefficient (Bouvier, 2015).

Ferrets are the gold standard animal model for IAV pathogenesis and transmission due to their susceptibility to human IAV strains, release of virus into the air within a range of aerosol sizes that can infect a susceptible host, similarity in lung physiology, infection kinetics and clinical symptoms when compared to humans (Belser, Eckert, Tumpey, & Maines, 2016; Belser, Pulit-Penaloza, & Maines, 2020; Lakdawala et al., 2015, 2011, 2013). In addition, ferrets first infected with IAV have been shown to be susceptible to Spn colonization and promote Spn airborne transmission (McCullers et al., 2010; Peltola, Boyd, McAuley, Rehg, & McCullers, 2006; Peltola, Rehg, & McCullers, 2004; Rowe, Karlsson, et al., 2019). Recently, Rowe and colleagues have demonstrated that the nasal microbiota promotes airborne transmission of H1N1pdm09 virus in the ferret model (Rowe et al., 2020). Together, these studies support a model of synergism between Spn and IAV regarding host virulence, Spn colonization, and Spn transmission. However, the benefit to IAV fitness in the host and airborne transmission has not been fully addressed in the context of coinfection, which may impact the spread of pandemic viruses such as the H1N1pdm09.

In this study, we examined the impact of Spn (serotype 2 strain D39 and serotype 19F strain BHN97) coinfection on airborne transmission of H1N1pdm09 in 6-months-old ferrets to address this gap in knowledge. Similar to previous studies, we observe that IAV-Spn coinfection results in enhanced bacterial burden and increased morbidity. Surprisingly, we found that coinfection with Spn reduced airborne transmission of H1N1pdm09 when compared to virus only infected controls. This reduction in transmission was not due to reduced stability within expelled droplets, which were tested at relevant indoor humidity and temperature conditions. We observed that coinfection with Spn reduced viral titers in the upper respiratory tract (URT) of coinfected animals compared to H1N1pdm09 only infected ferrets. Our data suggest that coinfection with Spn reduces the amount of virus expelled by infected ferrets thus reducing airborne transmission of naïve recipients and challenges the synergistic relationship between IAV and Spn. Rather, we propose an asymmetrical relationship where IAV infection increases Spn burden but Spn colonization reduces H1N1pdm09 URT replication and airborne transmission.

## Results

### Development of ferret model for IAV-Spn transmission

Only a limited number of studies have investigated IAV-Spn airborne transmission in the ferret model (McCullers et al., 2010; Rowe et al., 2020). To address this gap in knowledge, we utilized the ferret model to study the impact of Spn (serotype 2 strain D39) infection on airborne transmission of H1N1pdm09. Initially, a dose titration experiment was performed to identify the amount of Spn D39 required to colonize IAV infected ferrets. Ferrets were intranasally infected with H1N1pdm09 then two days later, close to the peak of viral shedding, Spn D39 was administered by the same route with doses of 10^5^, 10^6^ or 10^7^ colony forming units (CFUs) (Fig 1A). Nasal washes were collected on the experimentally infected donors every day for 14 days post-H1N1pdm09 infection (12 days post-Spn infection). Colonization of Spn is defined as detectable bacterial shedding in nasal washes for more than three days. Robust Spn colonization was observed in two donors, one given 10^5^ and the other given 10^7^ CFUs Spn (Fig 1B and 1D, respectively), whereas the 10^6^ CFUs Spn donor cleared the bacteria by 5 days post-infection (dpi) (Fig 1C). These data demonstrate that Spn can colonize ferrets after H1N1pdm09 infection at multiple doses.

**Figure 1:**
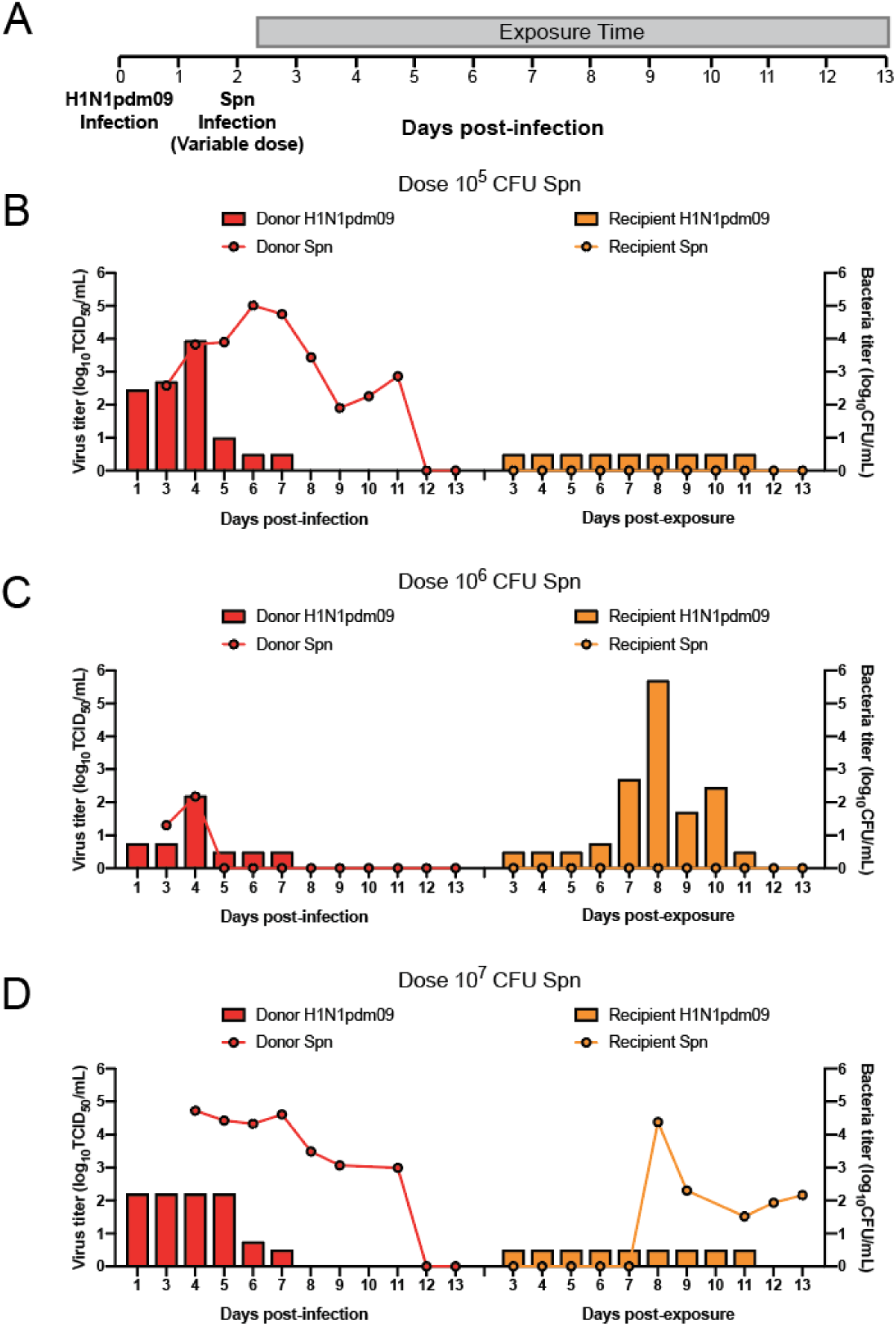
Ferrets as a model for airborne transmission during H1N1pdm09 and *Streptococcus pneumoniae* (Spn) D39 coinfections. (**A**) Experimental setup for the airborne transmission in ferrets. Donors (n=3) were inoculated with 10^6^ TCID_50_ H1N1pdm09 (A/CA/07/2009) and two days later with Spn (D39) with increasing doses between animals. Recipients (n=3) were placed in the transmission cages 6 hours post-bacterial inoculation. The gray box indicates the exposure time. Nasal washes were collected daily for up to 11 days and assayed for viral and bacterial titer. (**B, C, D**) Viral (bar graph) and bacterial (line graph) titers in donor (red) and recipient (orange) animals for each pair are indicated. The limit of detection for virus is 10^0.5^ TCID_50_ per mL and bacteria is 10^1^ CFU per mL. Days post-exposure is relative to time of viral inoculation.

To examine whether Spn and IAV can transmit through the air and establish an infection, we additionally performed an airborne transmission experiment. Six hours following D39 infection, naïve recipients were placed into the adjacent cage, which is separated from the donor animal by perforated metal plates. This prevents physical contact but allows for air flow from the donor animal to the recipient (Lakdawala et al., 2015, 2011). Recipients were exposed to the coinfected donors for eleven consecutive days with daily collection of nasal washes and assessment of clinical symptoms (Fig 1A). Transmission of both H1N1pdm09 and Spn is defined by detection of either pathogen in nasal secretions. Seroconversion for H1N1pdm09 and Spn on day 14 post-exposure was measured as a marker for exposure, but seroconversion alone was not counted as a transmission event. Airborne transmission of H1N1pdm09 was only observed in one pair, and transmission of Spn was not observed in this pair (Fig 1C). Spn only transmitted to one recipient who was paired with the donor that received 10^7^ CFU Spn. This animal displayed robust colonization, shedding Spn from 8 to 13 dpi, however, no detectable H1N1pdm09 was observed in the nasal secretions (Fig 1D). At the time of sacrifice (14 days post-exposure), respiratory tissue sections were collected and analyzed for presence of Spn, which was found in nasal turbinates (10^5^ CFU/mL) but no other anatomical site in this animal. We determine transmission efficiency based on shedding of virus or bacteria in nasal secretions. To confirm the transmission dynamics, we also quantified seroconversion to H1N1pdm09 and Spn (Table S1). Seroconversion matched bacterial and viral shedding patterns, except that the recipient who shed Spn also seroconverted for H1N1pdm09 but did not shed virus in nasal secretions (Fig 1D). These data indicate that at an inoculation dose of 10^7^ CFUs of Spn there was airborne transmission of Spn to the recipient. No other doses resulted in airborne transmission of Spn, thus this dose was selected for subsequent experiments.

### Coinfection with Spn D39 reduces transmission of H1N1pdm09

In the ferret model, H1N1pdm09 virus has consistently been reported to efficiently transmit to recipient animals through the air (Koster et al., 2012; Lakdawala et al., 2015, 2011; Maines et al., 2009; Paules et al., 2017; Pulit-Penaloza et al., 2018). Yet, in our initial study, transmission of H1N1pdm09 was observed in only one of three recipients, suggesting that the presence of Spn lowers virus transmissibility. To determine the impact of Spn on IAV transmission, we performed an H1N1pdm09 only group concurrently to ensure that the transmission of H1N1pdm09 was not dependent upon shifting the exposure window from 2 to 13 dpi, rather than classical exposure duration of 1 to 13 dpi (Lakdawala et al., 2015, 2011; Maines et al., 2009; Pulit-Penaloza et al., 2018). Six donors were infected with H1N1pdm09 and 2 days later three of these donors were infected with 10^7^ CFU of Spn D39. Recipients were placed into the transmission cage 6 hours after bacterial inoculation, and nasal secretion were assessed for Spn and H1N1pdm09 titers to determine transmission efficiencies. In accordance with previous H1N1pdm09 transmission studies, three of three of recipients exposed to H1N1pdm09 only infected donors shed virus in nasal secretions (Fig 2A). In contrast, coinfection with Spn decreased H1N1pdm09 transmission efficiency to one of three recipients (Fig 2B). Ferrets coinfected with H1N1pmd09 and Spn became colonized with Spn and shed detectable bacteria between 3 and 13 dpi (Fig 3B, red lines). Moreover, airborne transmission of Spn occurred in one of three recipient animals, with Spn shedding detected between 9 and 14 dpi (Fig 3B, orange lines). This is the same recipient that shed H1N1pdm09 (Fig 2B, orange bars), demonstrating airborne transmission of both pathogens to a single animal. Intriguingly, the temporal dynamics of transmission appear distinct for these two pathogens with shedding of H1N1pdm09 waning at day 10 post-exposure when bacterial shedding was beginning. Additionally, this recipient was still shedding Spn in nasal washes at the time of necropsy (16 dpi) (Fig 3B). Analysis of serum at 16dpi revealed antibodies specific to H1N1pdm09 but not Spn (Table S1), indicating there was insufficient time to mount a detectable immune response for Spn but not H1N1pdm09. The reduced transmission efficiency during coinfection strongly suggests that Spn coinfection decreases transmission of H1N1pdm09.

**Figure 2:**
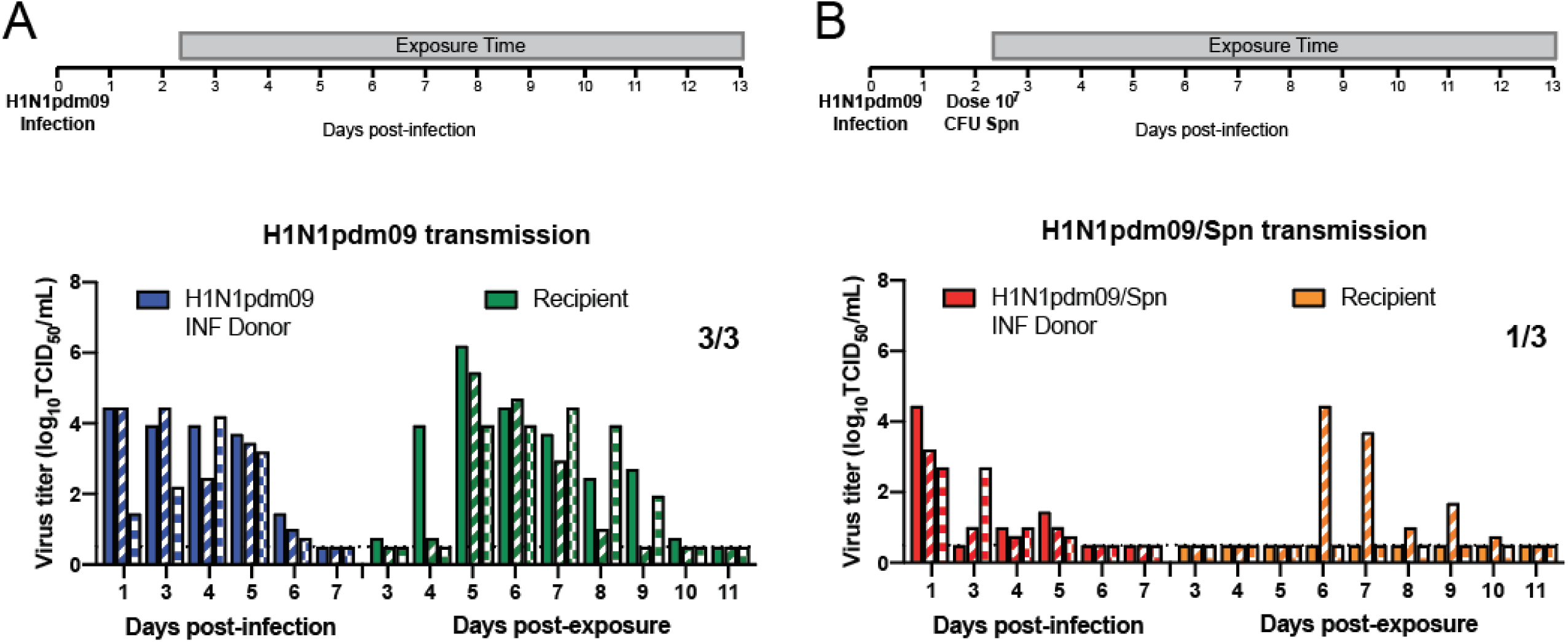
Airborne transmission of H1N1pdm09 is reduced upon coinfection with Spn (D39) in ferrets. Experimental setup for the airborne transmission in ferrets is shown above each panel. Donors (n=3) were infected with 10^6^ TCID_50_ H1N1pdm09 (A/CA/07/2009). Two days later, donors were not infected **(A)** or inoculated with 10^7^ CFUs of Spn (D39) **(B)**. Recipients were placed in the adjacent cage 6 hours later and exposed for 11 days (gray bar). Each bar represents an individual animal. Limit of detection is denoted by a dashed line. Days post-exposure is relative to time of viral inoculation.

**Figure 3:**
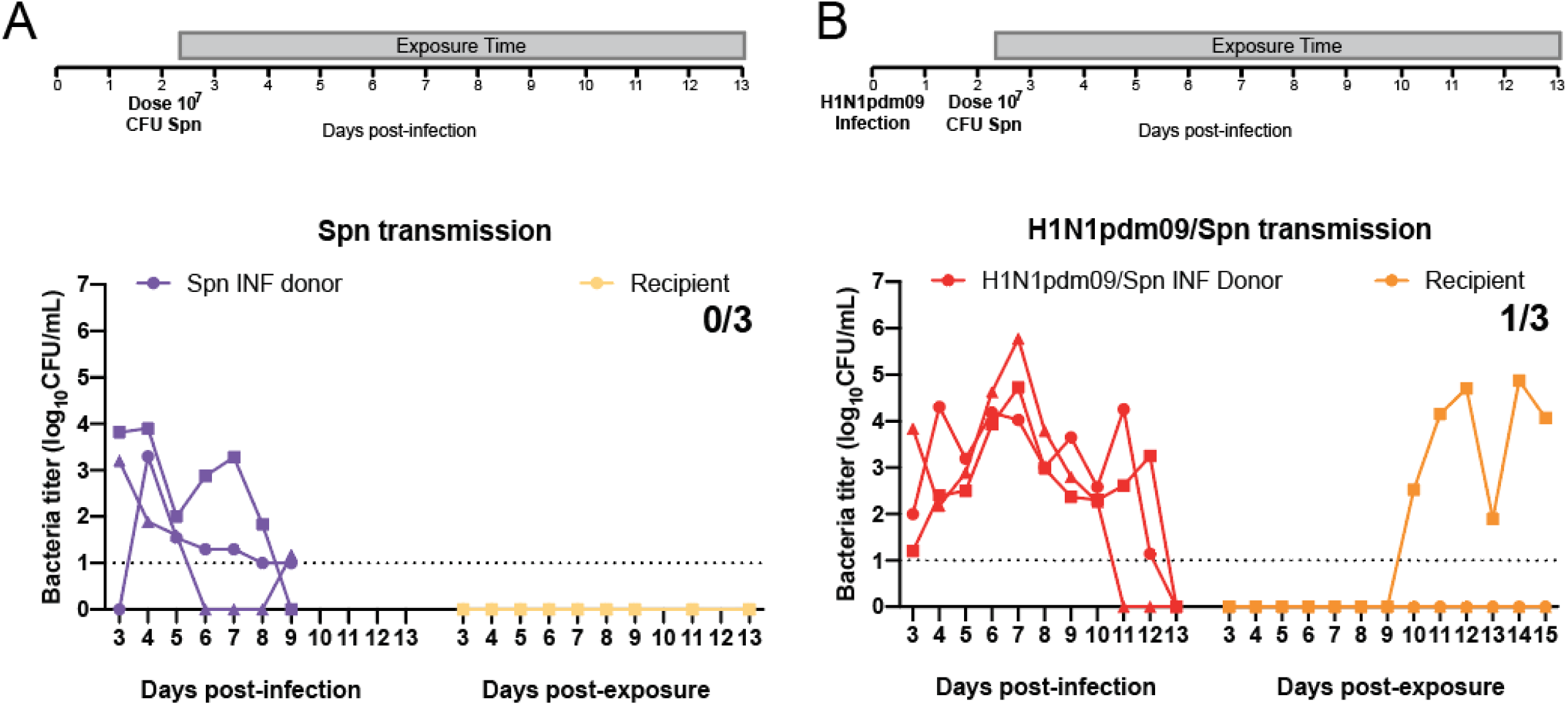
Spn D39 airborne transmission is not efficient in ferrets. Experimental setup for the airborne transmission in ferrets is shown above each panel. **(A)** Donors (n=3) were infected with 10^7^ CFUs of Spn (D39). **(B)** Donors (n=3) were infected with 10^6^ TCID_50_ H1N1pdm09 (A/CA/07/2009) and two days later, donors were inoculated with 10^7^ CFUs of Spn (D39). Recipients were placed in the adjacent cage 6 hours after bacterial infection and exposed for 11 days (gray bar). Bacterial titers from donors and recipient animals are indicated by each dot, which represents an individual animal. Limit of detection is denoted by a dashed line. For consistency, days post-exposure is relative to time of viral inoculation event for A where virus was not added.

### Inefficient airborne transmission of Spn D39 in the ferret model

To investigate whether H1N1pdm09 infection influenced Spn transmission, ferrets were inoculated with 10^7^ CFUs Spn D39 without prior IAV infection in a separate experiment (Fig 3A). Bacterial colonization occurred in the absence of H1N1pdm09 infection with the donors shedding over multiple days (Fig 3A, purple lines). While similar levels of Spn shedding were observed on days 1 and 2 post infection between Spn only and coinfected ferrets, the amount of shedding was reduced from three to seven dpi (Fig 6C). No transmission events were detected in the Spn only group (Fig 3A). In ferrets, the transmission of Spn appears to be low and necessitates prior IAV infection.

### Coinfection with Spn BHN97 reduces H1N1pdm09 transmission

Isolated in 1916, the model strain Spn D39 is a serotype 2 strain that is highly virulent in murine models of pneumococcal pathogenesis. Importantly, pathogenesis of Spn can be strain dependent in mice. For example, in contrast to D39, the clinical serotype 19F strain BHN97 colonizes to high density but does not typically cause invasive disease in mice (Rosch et al., 2014). In ferrets, previous studies have noted that the Spn strain can influence the synergism between these two pathogens (McCullers et al., 2010). To determine whether the reduction in IAV transmission was dependent upon Spn strain, we intranasally infected ferrets with H1N1pdm09 then two days later similarly administered 10^7^ CFUs of Spn BHN97 (Fig 4A) or mock infected with PBS (Fig 4C). Recipients were placed into the transmission cage 6 hours after bacterial inoculation, and nasal secretion were titered for Spn BHN97 and H1N1pdm09 to determine transmission efficiencies. Ferrets coinfected with H1N1pdm09 and BHN97 became colonized with Spn and shed bacteria as of 3 dpi (Fig 4B, red lines). However, two donor animals succumbed to the bacterial infection and were found dead in the cage on days 5 and 6 post-infection (Fig 4B, skull symbol) with high titers of BHN97 in all organs tested (Fig S3B and S3C, respectively). Blood collected at time of necropsy revealed the presence of Spn, demonstrating that these animals were septic.

**Figure 4:**
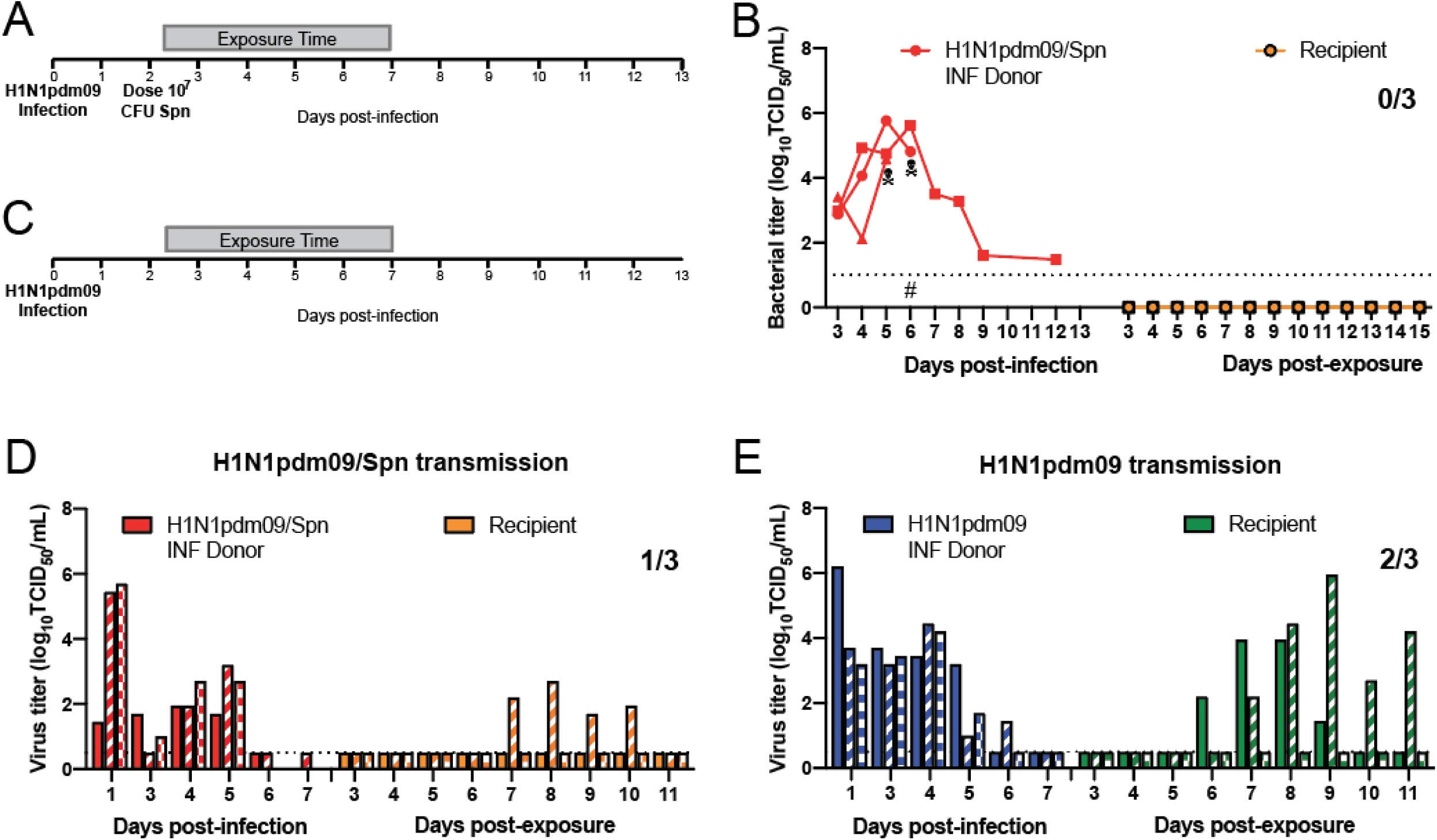
Spn BHN97 reduces airborne transmission of H1N1pdm09. In the experimental setup, donors (n=6) were infected with 10^6^ TCID_50_ H1N1pdm09 (A/CA/07/2009). Two days later, donors were either inoculated with 10^7^ CFUs of Spn (BHN97) (n=3) **(A)** or mock infected (n=3) **(C)**. Recipients were placed in the adjacent cage 6 hours after bacterial infection and exposed for 5 days (gray bar). **(B)** Bacterial titers from donors and recipient animals in panel A are indicated by each dot, which represents an individual animal. Skull symbol represents sudden death of a ferret and # the beginning of antibiotic treatment. H1N1pdm09 titers for coinfection **(D)** and IAV alone **(E)** where each bar represents an individual animal. Limits of detection for bacteria and IAV are denoted by a dashed line. Days post-exposure is relative to time of viral inoculation.

Based on the severe disease and death of these donor animals, the recipient animals for all pairs were separated such that the exposure time of all recipients was shortened to five days rather than 11 days. In this study, we did not observe any detectable bacterial shedding (Fig 4B, orange lines) or seroconversion (Table S1), suggesting that Spn BHN97 was not transmitted through the air to recipients. Coinfected donors shed H1N1pdm09 in nasal secretions till 5 dpi (Fig 4D, red bars), and H1N1pdm09 transmitted to one of three recipient animals (Fig 4D, orange bars). A concurrent H1N1pdm09 only transmission experiment was performed with the same five day exposure time as the coinfected group (Fig 4C), which resulted in donors shedding virus till 6 dpi and two of three recipients shedding virus from days six to eleven post-exposure (Fig 4E, green bars). Seroconversion for H1N1pdm09 was observed only in animals that also shed virus (Table S1). Thus, H1N1pdm09 airborne transmission was reduced from 66% to 33% in the presence of Spn. Given the low numbers of animals this difference is not significant but represents a trend toward a decrease in airborne H1N1pdm09 transmission to recipient animals. Taken together, these results suggest that the negative impact of Spn on H1N1pdm09 transmission is not strain specific.

### Stability of H1N1pdm09 in droplets is unaffected by the presence of Spn D39

Airborne transmission requires the pathogens to remain infectious in expelled droplets for prolonged periods of time (Tang, 2009). Previous work has suggested that Spn and IAV can physically interact with each other (Rowe, Meliopoulos, et al., 2019). Therefore, expelled droplets containing both Spn and IAV could impact the stability of the virus and consequently influence transmission. To determine whether the stability of H1N1pdm09 in expelled respiratory droplets decreased in the presence of Spn, we utilized a previously established stability assay (Kormuth et al., 2018, 2019). In this setup, viruses and Spn are mixed with airway surface liquid (ASL) collected from patient-derived human bronchial epithelial (HBE) cell cultures differentiated at an air-liquid interface to mimic physiologically relevant conditions. Briefly, small 1μL droplets were incubated for 2 hours at 43% relative humidity, which is representative of the environmental conditions during the ferret transmission experiments (Fig S1). The stability of H1N1pdm09 was determined in the presence and absence of Spn D39 with ASL from 4 different HBE cultures to account for patient variability (Fig 5A). The viral titers obtained from the droplets were compared to an equal volume of the virus, ASL, +/-Spn in a closed tube outside of the chamber (referred to as ‘no chamber’ control). The log_10_ decay for each HBE culture was calculated as the ratio of H1N1pdm09 titer in the no chamber control to the titer in droplets (Fig 5B). These data did not reveal a difference in H1N1pdm09 stability in the presence or absence of Spn (Fig 5A and 5B).

**Figure 5:**
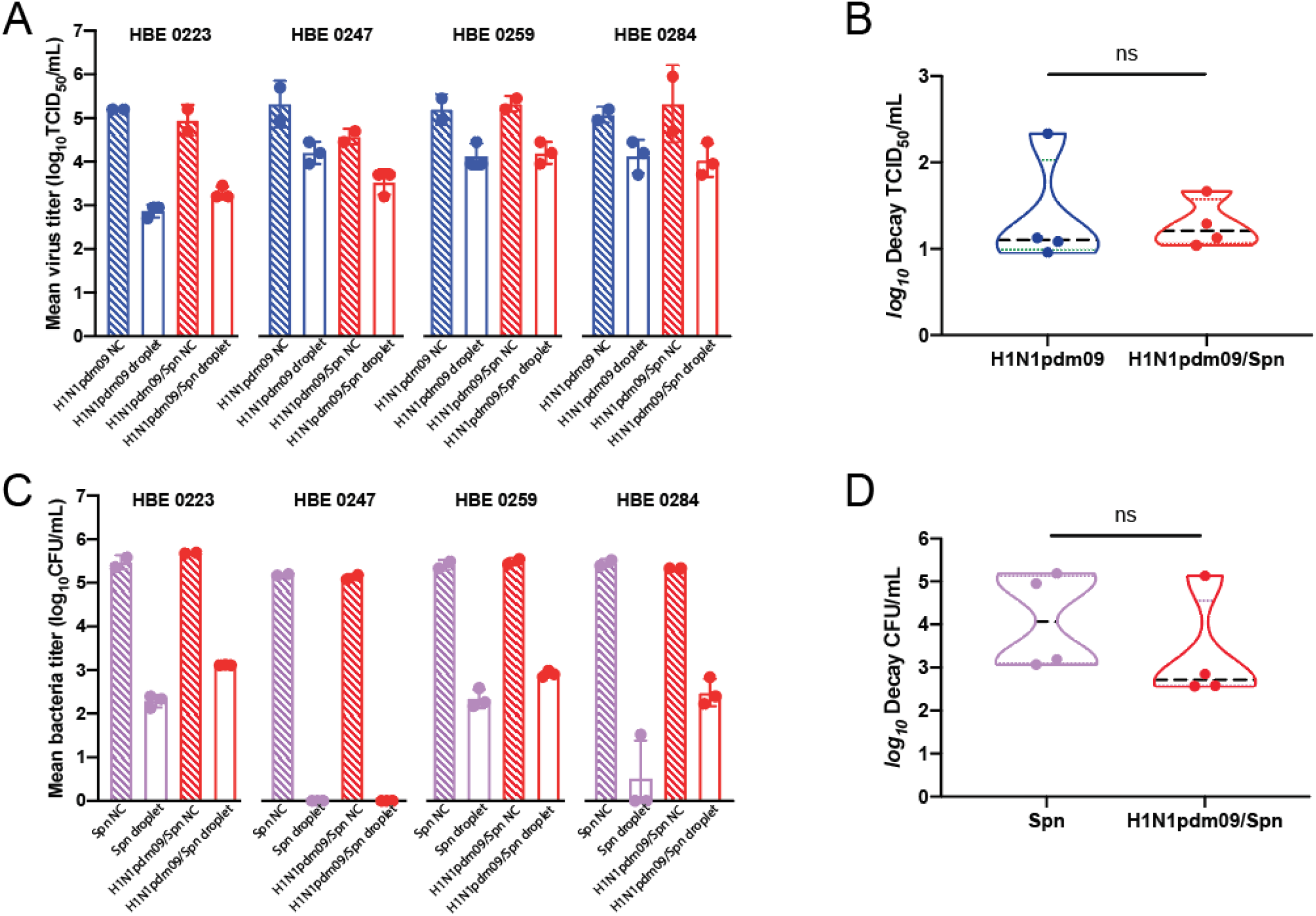
Stability of H1N1pdm09 in droplets is not affected by Spn D39. Airway surface liquid (ASL) from 4 different patient-derived human bronchial epithelial (HBE) cell lines was combined with H1N1pdm09 and/or Spn D39. Large 1μL droplets were aged for 2 hours at 43% relative humidity and normalized to 10μL bulk solution aged for 2 hours in closed tubes. **(A**,**C)** Raw TCID_50_/mL and CFU/mL for test and no chamber (NC) control samples. **(B**,**D)** Log_10_ Decay of H1N1pdm09 and Spn D39 after aging in ASL from different patient cell lines, with points representing biological replicates of unique patient-derived HBE ASLs. Significance was determined using Student’s *t*-test with Welch’s correction.

Conversely, to determine whether the presence of H1N1pdm09 contributed to increased stability of Spn in expelled droplets, Spn titers were assessed with and without H1N1pdm09 in ASL from 4 different HBE cultures (Fig 5C). The log decay of Spn titers in droplets compared to no chamber controls did not reveal a significant impact of H1N1pdm09 on Spn stability (Fig 5D). However, a trend toward increased stability in the presence of H1N1pmd09 was observed (Fig 5C and 5D). Interestingly, there were differences in the overall log decay between HBE cultures, where ASL from specific cultures produced a higher level of decay for both H1N1pdm09 (HBE 0223) and Spn (HBE 0247). Based on these data, stability of H1N1pdm09 in the presence of Spn does not correlate with the decreased transmission phenotype.

### Coinfection of H1N1pdm09 and Spn results in severe pathogenesis

Coinfection of Spn and IAVs results in a more severe disease outcome in humans, and mouse models of pneumococcal disease (McCullers & Rehg, 2002; Morens et al., 2008; Rudd et al., 2016). Consistent with these published findings, we found clinical symptoms were more severe in the coinfected donors as compared to those infected with H1N1pdm09 or Spn alone (Table S1), including scoring based on weight loss and activity (Fig S2A and S2B). A thick layer of nasal discharge was observed in animals coinfected with H1N1pdm09 and Spn, but not in animals with H1N1pdm09 or Spn alone (Fig S2C). Furthermore, coinfected ferrets displayed diarrhea, shallow breathing when put under anesthesia, and severe dehydration that required administration of subcutaneous fluids (Table S1). At 16 dpi, the recipient from Fig 2B and 3B, who was coinfected with both H1N1pdm09 and Spn, reached the 20% weight endpoint and was sacrificed. Dissemination of Spn was observed with high bacterial loads in all tissues tested, including the spleen and blood, demonstrating sepsis (Fig S2A). Coinfection with H1N1pmd09 and BHN97 caused two of three donors to rapidly succumb to bacterial sepsis on 5 and 6 dpi (Fig S3B and S3C). These findings are in accordance with previous studies that showed pneumococcal pathogenesis to be more severe when preceded by IAV (Kash et al., 2011; Mina et al., 2015; Peltola et al., 2006, 2004).

### Coinfection enhances bacterial titers while decreasing viral titers

Consistent with enhanced disease during coinfection, all coinfected ferrets had increased disease burden as compared to either viral or bacterial pathogen alone (Fig S2 and Table S1). Furthermore, Spn D39 titers were greater in the presence of H1N1pdm09 at later times post-infection with significant differences at 4 and 5 dpi (Fig 6C). Prior to Spn infection, viral titers in the nasal secretions were similar on day 1 post-H1N1pdm09 infection between the IAV only and coinfected groups (Fig 6A and 6B). The day following Spn infection, we observed that viral titers for H1N1pdm09 were significantly lower in nasal secretions upon secondary infection with Spn D39 (Fig 6A). Similarly, coinfection with Spn BHN97, resulted in lower H1N1pdm09 titers on 2 dpi but recovered to similar levels as the IAV only controls (Fig 6B). These data suggest that both Spn serotype 2 and 19F are inhibitory to H1N1pdm09 replication in the upper respiratory tract of ferrets, which may be associated with the reduced airborne transmission of H1N1pdm09 virus.

**Figure 6:**
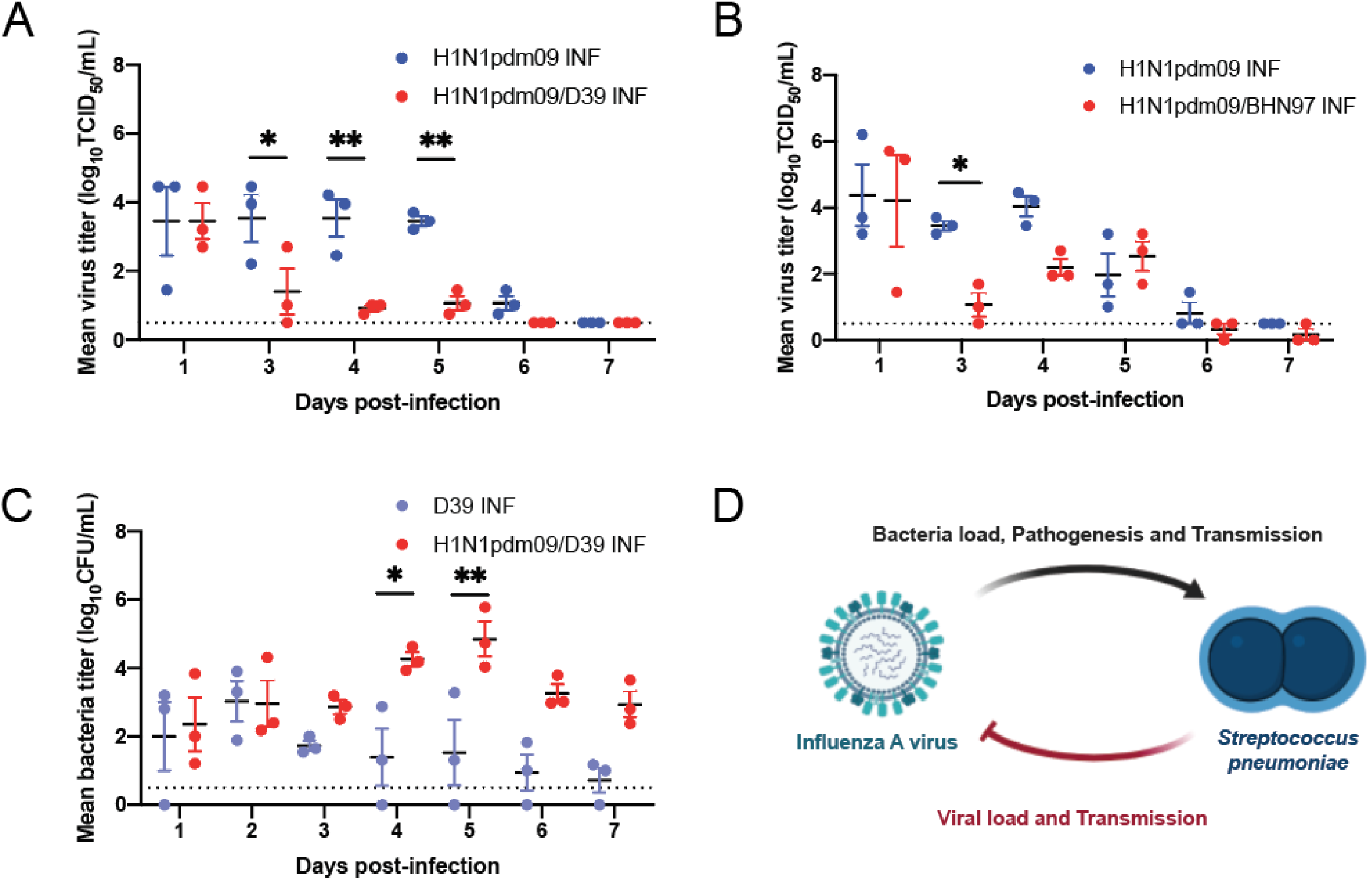
Coinfection in ferrets enhances bacterial titer while significantly reducing the viral titer. Comparison of mean virus titer of donors during the first week of infection between concurrent H1N1pdm09 alone versus coinfection with Spn D39 **(A)** or BHN97 **(B)**. Data represent the mean ± SEM for each group per day. Two-way ANOVA analysis was used to determine statistically significant differences (* p<0.05, ** p<0.005). (**C**) Comparison of mean bacterial titer of animals during Spn D39 infection and coinfection with preceding H1N1pdm09 infection. (**D**) Current model for interaction of IAV and Spn in the ferret model. H1N1pdm09 enhances the bacterial load, pathogenesis, and transmission in ferrets. The propagation of the bacteria in the host subsequently results in decreased viral load and airborne transmission of IAV. Schematic was created using BioRender.com.

## Discussion

Efficient airborne transmission is critical for the spread and public health burden of respiratory pathogens. Understanding how microbial communities contribute to the spread of pathogenic bacteria and viruses through the air is understudied. Based on the data presented here, we demonstrate that coinfection with Spn reduces H1N1pdm09 airborne transmission and replication in the URT. In contrast, coinfection of H1N1pdm09 promotes Spn airborne transmission, colonization of the URT, systemic dissemination, and pathogenesis. These data suggest an asymmetrical relationship between Spn and IAV, where IAV infection enhances Spn fitness within a host and between hosts but Spn infection negatively influences IAV replication and transmission fitness (Fig 6D).

Our current understanding of viral and bacterial coinfection reveals changes in disease pathogenesis, replication fitness within a host, and transmission between hosts, which can all influence the public health burden of respiratory pathogens. During the 2009 pandemic (H1N1pdm09), severe infections were attributed to secondary bacterial pneumonia, commonly caused by Spn (Macintyre et al., 2018; Palacios et al., 2009). The increased disease severity upon coinfection is strongly supported by animal models. Studies in mice and ferrets report increased morbidity and mortality in coinfected animals relative to single infection (Kash et al., 2011; Marks, Davidson, Knight, & Hakansson, 2013; McCullers et al., 2010; Mina et al., 2015; Peltola et al., 2006; Sharma-Chawla et al., 2016). Consistent with these previous data, we found that relative to a single infection, coinfected ferrets displayed enhanced disease as measured by increased weight loss, appearance of a thick layer of nasal discharge, diarrhea, shallow breathing when put under anesthesia, and severe dehydration that required administration of subcutaneous fluids. In contrast to previous studies with younger ferrets(Rowe et al., 2020), two of the ferrets coinfected with H1N1pdm09 and BHN97 rapidly succumbed and were found to have high titers of Spn in their blood and organs. Taken together these data highlight the severity of clinical outcomes during IAV and Spn coinfection in many relevant animal model systems.

Our study and others, in both mice and ferret models, have concluded that coinfection with IAV promotes Spn colonization (Marks et al., 2013; Rowe, Karlsson, et al., 2019; Wren et al., 2014, 2017). While we find that Spn colonization can occur in the absence of H1N1pdm09 infection, the bacterial titers are higher in the presence of the virus. Surprisingly, we observed that coinfection with Spn decreased H1N1pdm09 replication in URT as determined by viral titration of nasal washes. This observation is similar to recent data obtained from neonatal mice colonized with Spn prior to PR8 H1N1 infection (Ortigoza et al., 2018). The difference in enhancement of Spn colonization versus reduction in viral titers indicates an asymmetrical relationship between these two pathogens. Previous work has provided strong evidence that viruses from the URT of infected ferrets preferentially transmit through the air (Lakdawala et al., 2015; M. Richard et al., 2020). Therefore, the decrease in URT viral titers in the presence of Spn infection likely influences the transmissibility of the virus by reducing the amount of virus expelled into the environment.

Airborne transmission of IAV have been widely studied since the 2009 pandemic (Belser et al., 2020), yet there are relatively few studies on the role of coinfection in IAV transmission, partially due to limitations of mouse models to examine airborne transmission of IAV (Bouvier, 2015). Ferrets are the preferred animal model for IAV airborne transmission and have been used extensively by many groups to study transmission of H1N1pdm09 (Belser, Maines, Katz, & Tumpey, 2013). Consistent with our previous publications, we observed robust airborne transmission of H1N1pdm in ferrets, where 100% (3/3) or 66% (2/3) of naïve recipient ferrets in our control studies became infected with H1N1pdm09. The exposure time was slightly delayed compared to conventional transmission studies, since we began exposure 6 hours after Spn infection or 54 hours after H1N1pdm09 infection. Importantly, the H1N1pdm09 transmission study with only 66% transmission efficiency was performed with an exposure time of only 5 days (day 2-7 post-H1N1pdm09 infection) before termination due to lethality in the donor animals. This delay in exposure time may account for the transmission efficiency of less than 100%. Surprisingly, coinfection with Spn (either D39 or BHN97) led to a decrease in transmission to only 33% (2/6) of naïve recipients.

These changes between single and coinfections invite the question as to whether virus and bacteria can interact directly or, whether the positive effect of the virus on the bacteria and negative effect of the bacteria on the virus are mediated via changes in the host. Previous work captured a direct interaction between these pathogens (Rowe, Meliopoulos, et al., 2019). Specifically, Spn was shown to interact directly with whole-inactivated IAV and live IAV. Further, when pathogens were mixed, desiccated, and then rehydrated, the stability of the virus was enhanced by the presence of bacteria, specifically bacteria expressing a capsule. In this study, we investigate viral stability in the presence of airway surface liquid (ASL) isolated from human bronchial cells in a stability assay that is different than the one performed in Rowe et al. Under our conditions, we did not observe an impact of Spn on the stability of H1N1pdm09. Furthermore, we observed a lag in IAV and Spn shedding in the one coinfected recipient animal captured within our studies. Spn shedding was detected between 10 and 15 dpi, and the IAV shedding was detected between 6 and 10 dpi. The absence of substantial overlap in shedding of these pathogens, is consistent with a model that includes independent transmission of each pathogen. It is worth noting, that the presence of ASL influenced both pathogens in a sample-dependent manner. It is likely that the variation we observed across samples relates to mucus properties such as composition and presence of antimicrobial peptides. Finally, while our stability measurements did not detect measurable differences in IAV stability in the presence of Spn, we did detect a trend suggesting an influence of IAV on Spn stability. In half the samples, IAV increased the stability of Spn indicating that both IAV and ASL components may contribute to Spn stability in air droplets, and thus, are likely to influence Spn transmission efficiency.

A recent study by Rowe et al, examined the impact of nasal microflora on airborne transmission of H1N1pdm09 (Rowe et al., 2020). In this study, donor ferrets were treated with mupirocin ointment to reduce bacterial respiratory flora, then infected with H1N1pdm09. Antibiotic treated animals did not transmit the virus to naïve recipients, suggesting that nasal commensal bacteria are required for airborne transmission of IAV. In contrast to our studies, Rowe et al observed that inoculation with mupirocin resistant Spn BHN97, restored airborne transmission of H1N1pdm09. Thus, in a setup where animals have an altered nasal microbiome due to antibiotic treatment, Spn promoted IAV transmission. In contrast, we observed that in the presence of normal commensal flora Spn (strains D39 and BHN97) coinfection reduced airborne transmission of H1N1pdm09. It is possible that variations in nasal flora due to antibiotic treatment may explain these differences in transmission phenotypes. These data point towards a complex relationship between microbial communities and airborne transmission of IAV.

There are numerable ways the microbiota could influence attachment and entry of IAV into host cells, IAV replication, maturation of viruses with host cells, egress and dissemination, and interaction between coinfecting viral and bacterial pathogens (Jones, Le Sage, & Lakdawala, 2020). It is already clear that changes in glycosylation of the host incurred during IAV infection influence subsequent Spn colonization, and similarly changes in glycosylation via Spn affect subsequent IAV infection (Ortigoza et al., 2018; Siegel et al., 2014). It is likely that changes in host glycosylation by Spn and other bacteria also influence entry and egress of IAV, as well as its distribution in tissues across the respiratory tract. Similarly, the host immune status will vary based on exposure, and may differ dramatically depending on resident microbiota and dynamics of infections for each pathogen in a coinfection. Most animal models of viral and bacterial coinfections do not account for variations in ASL components and microbiota. Combined, our study and recent studies of others in IAV-Spn coinfections propose that these variables play critical roles in the airborne transmission of respiratory pathogens.

## Materials and methods

### Cells, viruses, and bacteria

MDCK (Madin-Darby canine kidney, obtained from ATCC) were grown at 37°C in 5% CO_2_ in MEM medium (Sigma) containing 5% Fetal Bovine Serum (FBS, HyClone), penicillin/streptomycin and L-glutamine. Reverse genetics plasmids of A/California/07/2009 were a generous gift from Dr. Jesse Bloom (Fred Hutch Cancer Research Center, Seattle) and were rescued as previously described in (Lakdawala et al., 2011). The viral titers were determined by tissue culture infectious dose 50 (TCID_50_) using the endpoint titration method on MDCK cells for H1N1pdm09 (Pulit-Penaloza et al., 2018). The bacterial strains used in this study were graciously provided by Dr. Hasan Yesilkaya (wild type Spn serotype 2 strain D39) (Andrew et al., 2018) and Dr. Jason Rosch (wild type Spn serotype 19F strain BHN97) (Rowe, Karlsson, et al., 2019). Bacteria were grown from frozen stocks by streaking on TSA-II agar plates supplemented with 5% sheep blood (BD). Cultures were generated in fresh Columbia broth (Thermo Fisher) and incubated at 37°C and 5% CO_2_ without shaking. Pneumococci in nasal washes and tissues were determined by plating serial dilutions onto blood agar plates incubated at 37°C overnight.

### Animal ethics statement

Ferret transmission experiments were conducted at the University of Pittsburgh in compliance with the guidelines of the Institutional Animal Care and Use Committee (approved protocol #19075697). Humane endpoints for this study included body weight loss exceeding 20% (relative to weight at challenge) and a prolonged clinical activity score of 3 based on the system designed by Reuman et al ^42^. Animals were sedated with approved methods for all procedures. Isoflurane was used for all nasal wash and survival blood draw, ketamine and xylazine were used for sedation for all terminal procedures followed by cardiac administration of euthanasia solution. Approved University of Pittsburgh DLAR staff administered euthanasia at time of sacrifice. After the first ferret rapidly succumbed to the H1N1pdm09-BHN97 coinfection and was found to have bacteria disseminated to all body sites, the remaining coinfected ferrets were subcutaneously administered 5 mg/kg Baytril, twice a day for 7 days.

### Animals

Five to six months old male ferrets were purchased from Triple F Farms (Sayre, PA, USA). All ferrets were screened for antibodies against circulating influenza A and B viruses, as determined by hemagglutinin inhibition assay (as described in(Lakdawala et al., 2011)) using the following antigens obtained through the International Reagent Resource, Influenza Division, WHO Collaborating Center for Surveillance, Epidemiology and Control of Influenza, Centers for Disease Control and Prevention, Atlanta, GA, USA: 2018-2019 WHO Antigen, Influenza A (H3) Control Antigen (A/Singapore/INFIMH-16-0019/2016), BPL-Inactivated, FR-1606; 2014-2015 WHO Antigen, Influenza A(H1N1)pdm09 Control Antigen (A/California/07/2009 NYMC X-179A), BPL-Inactivated, FR-1184; 2018-2019 WHO Antigen, Influenza B Control Antigen, Victoria Lineage (B/Colorado/06/2017), BPL-Inactivated, FR-1607; 2015-2016 WHO Antigen, Influenza B Control Antigen, Yamagata Lineage (B/Phuket/3073/2013), BPL-Inactivated, FR-1403. Clinical symptoms such as weight loss, temperature and Reumann activity score (Reuman, Keely, & Schiff, 1989) were recorded during each nasal wash procedure and other symptoms such as sneezing, coughing, lethargy or nasal discharge were noted during any handling events. Animals were given A/D diet twice a day to entice eating once they reached 10% weight loss. A summary of clinical symptoms for each study are provided in Supplemental Table 1.

Tissues and blood were collected to assess viral and bacterial titers. Tissues were weighed and homogenized in sterile PBS at 5% (nasal turbinate, spleen, and trachea) or 10% (lung) weight per volume. The soft palate was homogenized in 1 mL PBS. Bacterial titers were assessed by plating serial dilutions on blood agar plates.

### Transmission studies

Our transmission caging setup is a modified Allentown ferret and rabbit bioisolator cage similar to those used in (Lakdawala et al., 2015, 2011). For each study, three donor ferrets were anesthetized by isofluorane and inoculated intranasally with 10^6^ TCID_50_/500uL of A/California/07/2009. For coinfection experiments, the IAV-infected donors were inoculated with 10^5^, 10^6^ or 10^7^ CFU/500uL of *Streptococcus pneumoniae* D39 at 48 h post-infection. At 6 hours post-bacterial infection, a recipient ferret was placed into the adjacent cage, which is separated by two staggered perforated metal plates welded together one inch apart. Nasal washes were collected from each donor and recipient every day for 13 days. To prevent accidental contact or fomite transmission by investigators, the recipient ferret was handled first and extensive cleaning of all chambers, biosafety cabinet, and temperature monitoring wands was performed between each recipient and donor animal and between each pair of animals. Sera was collected from donors and recipients upon completion of experiments to confirm seroconversion. Environmental conditions were monitored every hour using a HOBO UX100-011 data logger (Onset) and ranged between 20-22°C with 44-50% relative humidity (Fig S1). To ensure no accidental exposure during husbandry procedures, recipient animal sections of the cage were cleaned first then then infected side, three people participated in each husbandry event to ensure that a clean pair of hands handled bedding and food changes. One cage was done at a time and a 10 min wait time to remove contaminated air was observed before moving to the next cage. New scrappers, gloves, and sleeve covers were used on subsequent cage cleaning.

### Serology assay

Analysis of neutralizing antibodies from ferret sera was performed as previously described(Lakdawala et al., 2011). Briefly, the miconeutralization assay was performed using 10^3.3^ TCID_50_ of A/California/07/2009 virus incubated with 2-fold serial dilutions of heat-inactivated ferret sera. The neutralizing titer was defined as the reciprocal of the highest dilution of serum required to completely neutralize infectivity of 10^3.3^ TCID_50_ of virus on MDCK cells. The concentration of antibody required to neutralize 100 TCID_50_ of virus was calculated based on the neutralizing titer dilution divided by the initial dilution factor, multiplied by the antibody concentration.

### Spn ELISA

96-well ELISA plates (Immulon) were coated with 50 μL of 5×10^7^ CFU/mL of Spn in coating buffer (KPL) and incubated overnight at 4°C. Plates were blocked for 1 h at room temperature with 150 μL PBS with 0.01% Tween-20, 3% normal goat serum and 0.5% milk powder. Ferret sera was heat inactivated at 56°C for 30 minutes. Two-fold serial dilutions were performed in a blocking buffer and incubated on the ELISA plate for 2 hours at room temperature. After washing with PBS-0.01% Tween-20, plates were incubated with peroxidase-conjugated goat anti-ferret IgG (Jackson). SureBlue TMB peroxidase substrate (KPL) was added to each well and the reaction stopped with 250 mM HCl. Absorbance was read at 450 nm.

### Stability experiments

HBE cultures were obtained from the University of Pittsburgh Airway Tissue Core through an IRB-approved protocol from excess human lung tissue from transplantation. ASL was acquired by combining PBS washes of HBEs. H1N1pdm09 was prepared in antibiotic-free media. H1N1pdm09 was combined with ASL with and without D39. Alternately, D39 was combined with ASL with and without H1N1pdm09 and 10×1μL droplets were placed, in triplicate, into small 4-well plates. Droplets were then exposed to 43% relative humidity in a chamber that had been conditioned using a saturated salt solution. Chambers containing a temperature and humidity logger were closed and placed in a biosafety cabinet for 2 hours. To control for differences in virus or bacteria present in stock solutions, 10μL of bulk solution was aged for 2 hours, in duplicate, in closed Eppendorf tubes. Following aging, virus and bacteria were resuspended in 500μL of PBS. Bacteria was titered immediately on sheep’s blood agar plates. Samples were then frozen and titered later using MDCKs. Log_10_ decay was calculated as previously described (Yang, Elankumaran, & Marr, 2012). Log_10_ Decay= log_10_(NC control/Aged experimental sample).

## Acknowledgments

This work was supported by the National Institute of Allergy and Infectious Diseases (HHSN272201400007C (Rothman PI) SSL (subcontractor)); the NIH and the Cystic Fibrosis Foundation Research Development Program (grant P30DK072506 to the University of Pittsburgh, SSL); and the NIH 1R01AI139077-01A1 (NLH). Additional funding for SSL includes the American Lung Association Biomedical Research grant. We are also grateful for additional support from the Eberly Family Trust (NLH).

## Author Contributions

KMB, VL, NLH and SSL designed the experiments, analyzed, and interpreted the data, and wrote the manuscript. KMB, VL, AJF, JEJ, GHP and AJA performed the experiments. All authors edited and approved the manuscript.

---

**Figure S1:**
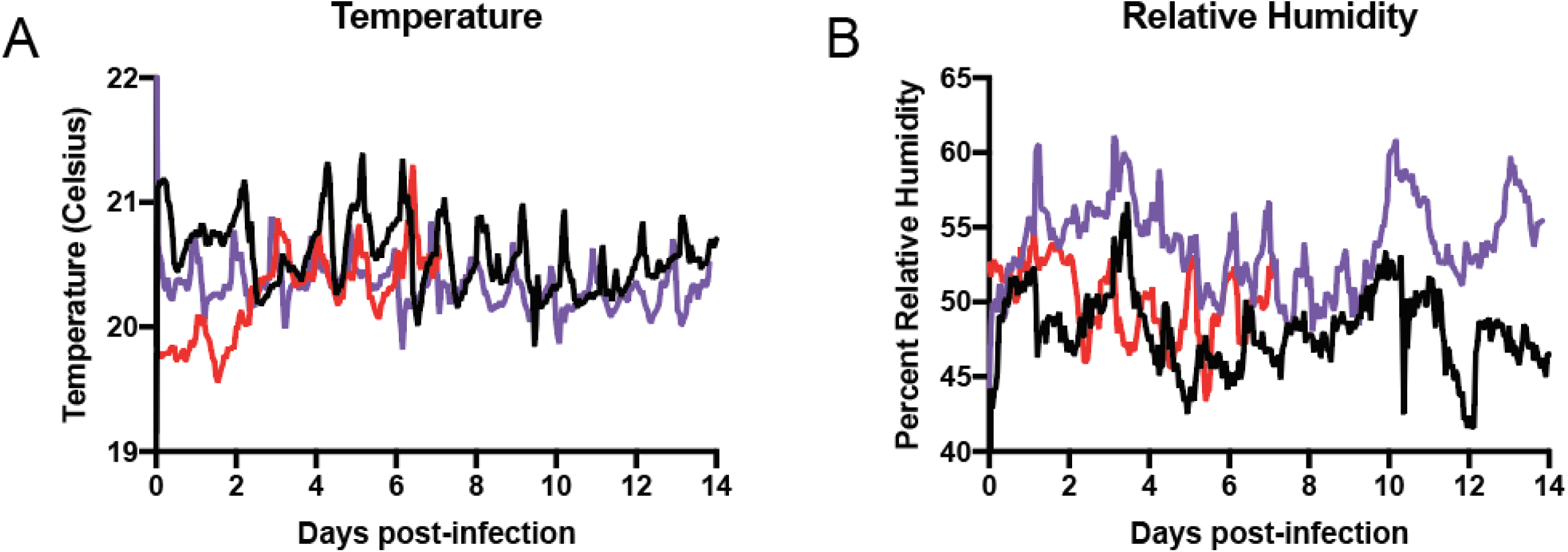
Temperature and relative humidity during transmission studies. Temperature **(A)** and relative humidity **(B)** for the coinfection and H1N1pdm09 only transmission studies, which were performed concurrently (D39; black line and BHN97; red line) while Spn only transmission (purple line) was performed separately.

**Figure S2:**
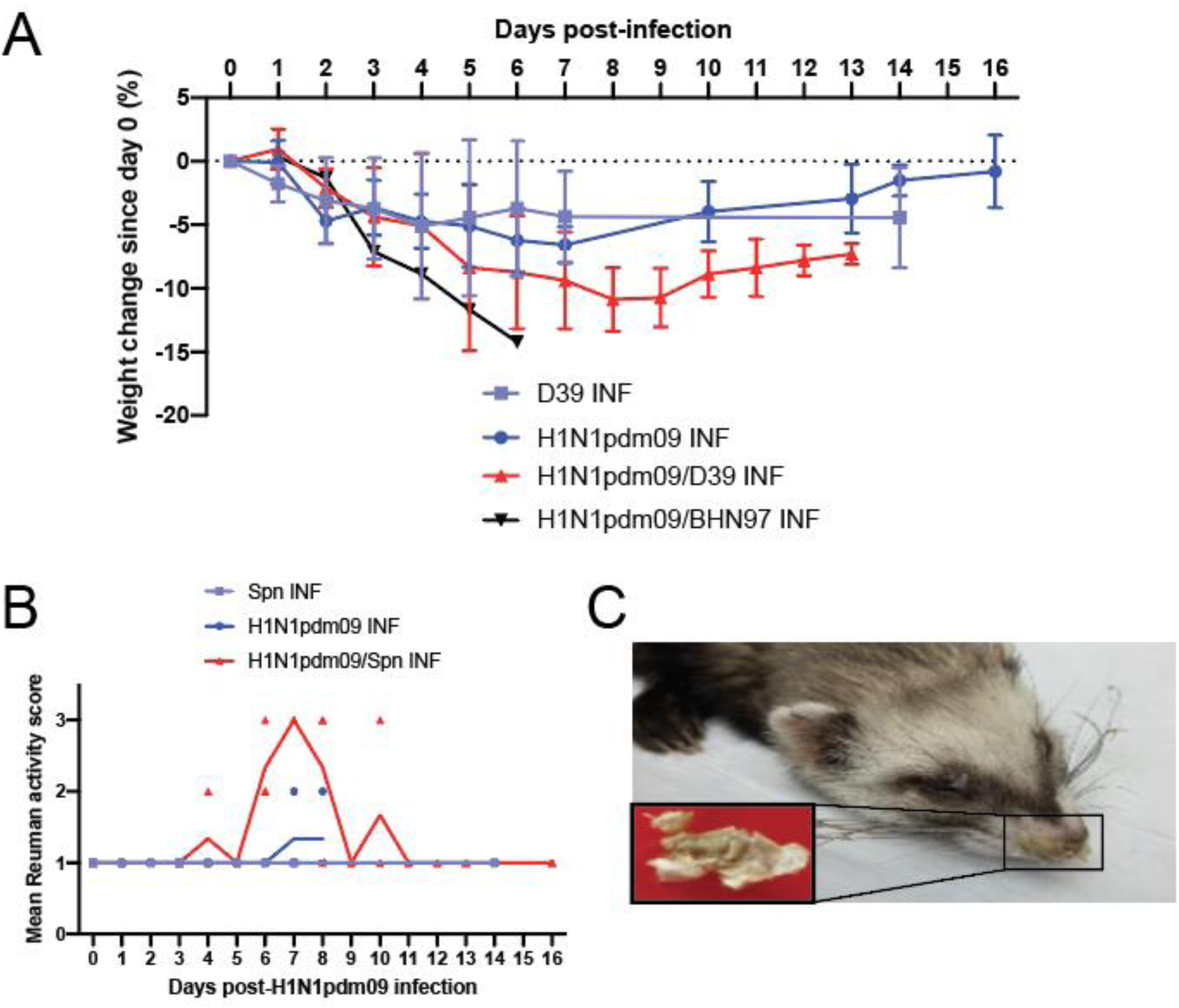
H1N1pdm09 and Spn coinfection causes more severe pathogenesis. **(A)** Comparison of weight loss from infected (INF) donors. **(B)** Reuman activity score of donor ferrets infected with Spn alone, H1N1pdm09 alone or coinfected with both pathogens at the indicated days post-infection. A score of 1 is fully alert and playful, 2 is alert but not playful and 3 is neither alert nor playful. **(C)** Coinfected ferrets developed a crusty nose and nasal discharge on whiskers. Extremely thick mucus was removed upon sedation (inset) and contained Spn upon streaking on blood agar plate (not shown).

**Figure S3:**
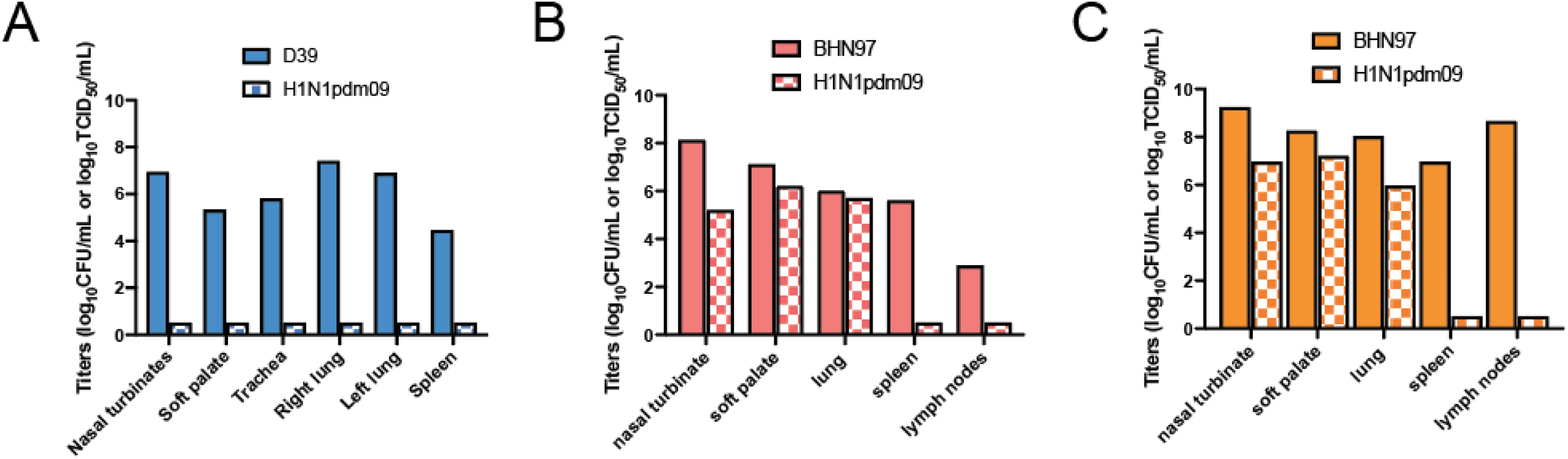
H1N1pdm09 and Spn titers from coinfection ferrets at endpoint. **(A)** D39 and H1N1pdm09 titers for the coinfected recipient from Fig 2B and 3B. Titers of BHN97 (log_10_ CFU/mL) and H1N1pdm09 (log_10_ TCID_50_/mL) from coinfected donors that succumb to bacterial infection of day 5 **(B)** and 6 **(C)** post-infection.

**Table S1:** Summary of clinal signs in donor and recipient ferrets

| Figure | Dose | | Status | Transmission efficiency | | Antibody titers\$ | | Clinical signs | | | | |
| --- | --- | --- | --- | --- | --- | --- | --- | --- | --- | --- | --- | --- |
|  | H1N1pdm09 | Spn |  | H1N1pdm09 | Spn | H1N1pdm09 | Spn | Temperature increase* | Weight loss >10% | max Reuman activity score<sup>+</sup> | Sneezing or coughing | Other clinical symptoms |
| 1B | 10<sup>6</sup> | 10<sup>5</sup> | Donor |  |  | 3620 | 150 | 0/1 | 1/1 | 3 | 1/1 | hardened nasal discharge, dehydration |
|  |  |  | Recipient | 0/1 | 0/1 | <20 | <100 | 0/1 | 0/1 | 1 | 0/1 |  |
| 1C | 10<sup>6</sup> | 10<sup>6</sup> | Donor |  |  | 2560 | <100 | 1/1 | 1/1 | 1 | 0/1 |  |
|  |  |  | Recipient | 1/1 | 0/1 | 2560 | 100 | 0/1 | 0/1 | 1 | 0/1 |  |
| 1D | 10<sup>6</sup> | 10<sup>7</sup> | Donor |  |  | 3225 | 1200 | 1/1 | 1/1 | 3 | 1/1 | hardened nasal discharge, dehydration |
|  |  |  | Recipient | 0/1 | 1/1 | 160 | 1200 | 1/1 | 0/1 | 3 | 1/1 | hardened nasal discharge, dehydration |
| 2A | 10<sup>6</sup> | NA | Donor |  |  | 1280, 1280, 1280 | <100, <100, <100 | 1/3 | 0/3 | 2, 1, 1 | 0/3 |  |
|  |  |  | Recipient | 3/3 |  | 640, 640, 1613 | <100, <100, <100 | 2/3 | 3/3 | 3, 2, 3 | 0/3 |  |
| 2B/3B | 10<sup>6</sup> | 10<sup>7</sup> | Donor |  |  | 1810, 1613, 1613 | 6400, 400, 1200 | 2/3 | 2/3 | 3, 3, 3 | 3/3 | hardened nasal discharge, dehydration (3/3) diarrhea (2/3) |
|  |  |  | Recipient | 1/3 | 1/3 | <20, 1613, <20 | <100, <100, <100 | 0/3 | 1/3 | 2, 3, 1 | 1/3 | hardened nasal discharge, dehydration, diarrhea, sepsis (1/3) |
| 3A | NA | 10<sup>7</sup> | Donor |  |  | ND | 75, 200, 400 | 2/3 | 1/3 | 1, 1, 1 | 0/3 |  |
|  |  |  | Recipient |  | 0/3 | ND | <100, <100, <100 | 0/3 | 0/3 | 1, 1, 1 | 0/3 |  |
| 4B/4D | 10<sup>6</sup> | 10<sup>7</sup> | Donor |  |  | ND, 80, ND | ND, 800, ND | 3/3 | 3/3 | 3, 3, 3 | 0/3 | hardened nasal discharge, dehydration (3/3) |
|  |  |  | Recipient | 1/3 | 0/3 | <20, 80, <20 | <200, <200, <200 | 0/3 | 0/3 | 1, 1, 1 | 0/3 | dehydration (1/3) |
| 4E | 10<sup>6</sup> | NA | Donor |  |  | 80, 113, 160 | ND | 0/3 | 1/3 | 2, 1, 1 | 0/3 |  |
|  |  |  | Recipient | 2/3 |  | 226, 16, <20 | ND | 0/3 | 1/3 | 1, 1, 1 | 0/3 |  |

\$ H1N1pdm09 antibody titers were determined by microneutralizations and Spn antibody titers were determined by whole bacteria ELISA
\* Temperature increase is defined as >1.5deg from day 0 temperature. Weight loss determined as > 10% of day 0 weight
<sup>+</sup> Maximum Reuman activity score. Number of animals with a score of >1 for more than 2 consecutive days (max score). 1-fully playful, 2-alert but not playful, 3-neither alert nor playful
ND = Not determined; NA = Not applicable

